# Acidic microenvironment shaped by lactate accumulation promotes pluripotency through multiple mechanisms

**DOI:** 10.1101/282475

**Authors:** Wen-Ting Guo, Shao-Hua Wang, Xiao-Shan Zhang, Ming Shi, Jing Hao, Xi-Wen Wang, Kai-Li Gu, Fei-Fei Duan, Ying Yan, Xi Yang, Chao Zhang, Le-Qi Liao, Yangming Wang

## Abstract

Enhanced glycolysis is a distinct feature associated with numerous stem cells and cancer cells. However, little is known about its regulatory roles in gene expression and cell fate determination. Here we show that acidic environment shaped by lactate accumulation promotes the self-renewal and pluripotency of both mouse and human embryonic stem cells (ESCs). Mechanistically, acidic pH reduces the tri-methylation of H3K27 globally at transcriptional start sites to partially prevent ESC differentiation. In addition, acidic pH stabilizes a large number of mRNAs including pluripotency genes. Furthermore, we found that AGO1 protein is downregulated at acidic conditions, leading to the de-repression of a subset of microRNA targets in low-pH treated ESCs. Altogether, our study provides insights into mechanisms whereby acidic microenvironment produced by enhanced glycolysis regulates gene expression to determine cell fate and has broad implications in the fields of regenerative medicine and cancer biology.

## Introduction

All cells require energy to maintain a high degree of order for survival and physiological functions. For animals the energy is derived from food molecules by well-organized catalytic processes degrading organic molecules such as proteins, lipids and polysaccharides. Two fundamental metabolic processes generating energy are glycolysis and oxidative phosphorylation (OXOPHOS). While glycolysis generates only 2 ATP molecules per glucose, OXOPHOS generates 38 ATP molecules per glucose. During OXOPHOS glucose is completely oxidized into CO_2_ and H_2_O. In contrast, glycolysis provides intermediate metabolites for various anabolic processes to synthesize nucleotides, lipids and amino acids^1, 2^. Interestingly, previous studies have shown that cancer cells and pluripotent stem cells (PSCs) prefer glycolysis rather than more efficient OXOPHOS for energy supply even in the presence of sufficient oxygen^3–5^. Likewise, during reprogramming, the acquisition of pluripotency is associated with the shift from oxidative to glycolytic metabolism. In contrast, during ESC differentiation, decreased glycolysis is observed with the energy supply of cells becoming increasingly dependent on OXOPHOS^6, 7^. In unicellular organisms, the metabolism switch is made possible simply by responding to the abundance of nutrient supplies in the environment. However, due to sufficient and constant nutrient supplies at most time, the metabolism switch of cells in multiple cellular organisms like mammals is regulated by sophisticated genetic circuits. Specifically in PSCs, various pluripotency-associated genes including Myc^8, 9^, Esrrb and Zic3^10^, and miRNAs including miR-290 and miR-200 families^11, 12^ have been shown to regulate their preference for glycolysis over OXOPHOS. Additionally, Hif1a is shown to promote the switch from bivalent to exclusively glycolytic metabolism during ESC to EpiSC transition^13^. These findings strongly suggest the importance of metabolism regulation in PSCs. However, the functional role and the working mechanism of different metabolic pathways in cell fate determination are still at large.

PSCs can undergo rapid self-renewal while retaining the ability to differentiate into all somatic cells in the body. Understandably, rapid proliferation requires the synthesis of a large quantity of biomolecules. For this reason, glycolysis has been proposed to provide abundant anabolic intermediates to support rapid proliferation^1, 2^. Furthermore, recent studies demonstrate that glycolysis play functional roles in keeping PSCs undifferentiated^9, 14^. The exact mechanism by glycolysis to promote self-renewal and pluripotency is not well understood. Chromatin modification is one of the most important processes controlling the self-renewal and pluripotency of PSCs^15, 16^. Chromatin modifiers require various cofactors and substrates for the chemical reaction of modification processes. Recent studies found that alpha-ketoglutarate and acetate generated from glucose flux are essential for histone methylation and acetylation by acting as a cofactor for histone demethylases^17^ and a substrate for histone acetyltransferases^14^, respectively. These studies reveal important functions of intermediate metabolites from glycolysis in regulating epigenetic modifications in PSCs. Nevertheless, besides many intermediate metabolites, glycolysis also produces a large amount of lactate which is transported into the extracellular space of PSCs. In fact, PSCs convert more than half the glucose consumed to lactate in vivo and in vitro^9, 18, 19^. It is estimated that the concentration of lactate can reach 16 mM and 100 mM in mouse and human blastocyst over a 24-hr period^20^, respectively. Hence, aerobic glycolysis can generate a very high concentration of lactate to alter the microenvironment of PSCs in vivo. Functionally, acidic microenvironment has been reported to enhance invasive and angiogenic potential of tumor cells^21–23^. However, the function of acidic pH has not been thoroughly investigated in PSCs. Recently, through inhibiting Na^+^-H^+^ exchanger DNhe2 (fly) or NHE1 (mouse), *Ulmschneider et al* found that the increase of intracellular pH is important for the differentiation of adult fly follicle stem cells and mouse ESCs^24^, implicating the function of lactic acid in regulating the self-renewal and pluripotency of stem cells.

In this study, we show that acidic microenvironment shaped by lactate accumulation play important functions in sustaining the self-renewal and pluripotency of PSCs. Lowering pH of culture media below 7 is sufficient to maintain the self-renewal and pluripotency of both mouse and human ESCs in normally differentiation conditions. Mouse ESCs can be maintained at pluripotent state for long term in basal media containing only LIF at low pH, replacing the requirement for chemical inhibitors to GSK3 and MEK. Mechanistically, we showed that low pH worked through reducing the trimethylation of H3K27 at transcriptional start sites, stabilizing mRNAs of pluripotency genes and suppressing miRNA activities. Together, these findings significantly improve our understanding of how acidic microenvironment produced by enhanced glycolysis regulates gene expression to control cell fate and have broad implications for the fields of regenerative medicine and cancer biology.

## Results

### Lactic acid but not lactate promotes the self-renewal and pluripotency of mouse ESCs

To monitor ESC differentiation process, we constructed mESCs expressing d2EGFP under the control of Rex1 promoter. The expression of GFP correlates with the activity of Rex1 promoter and therefore the pluripotency state of ESCs^25^. Rex1-d2EGFP ESCs expressed high levels of GFP when cultured in 2i+LIF media. However, they almost completely lost the expression of GFP when cultured in basal media without 2i and LIF for around 2 days (**Fig. 1a, b**). To investigate the influence of lactate generated by glycolysis on the pluripotency of ESCs, we supplemented basal media with 10 mM lactic acid or lactate salt. Surprisingly, lactic acid but not lactate sustained high levels of Rex1-d2EGFP expression (**Fig. 1a, b**). These data suggest that extracellular acidity caused by glycolysis is sufficient to support the pluripotency state of mouse ESCs for at least a short period. The addition of 10 mM lactic acid lowers the pH of basal media from ~7.4 to ~6.8. Consistent with acidic environment but not lactate salt promoting the self-renewal and pluripotency of ESCs, adjusting pH to 6.8 by other inorganic acids, including hydrochloride acid and sulphuric acid, also blocked the loss of Rex1-d2EGFP expression in the absence of 2i and LIF (**Fig. 1c**).

**Figure 1.**
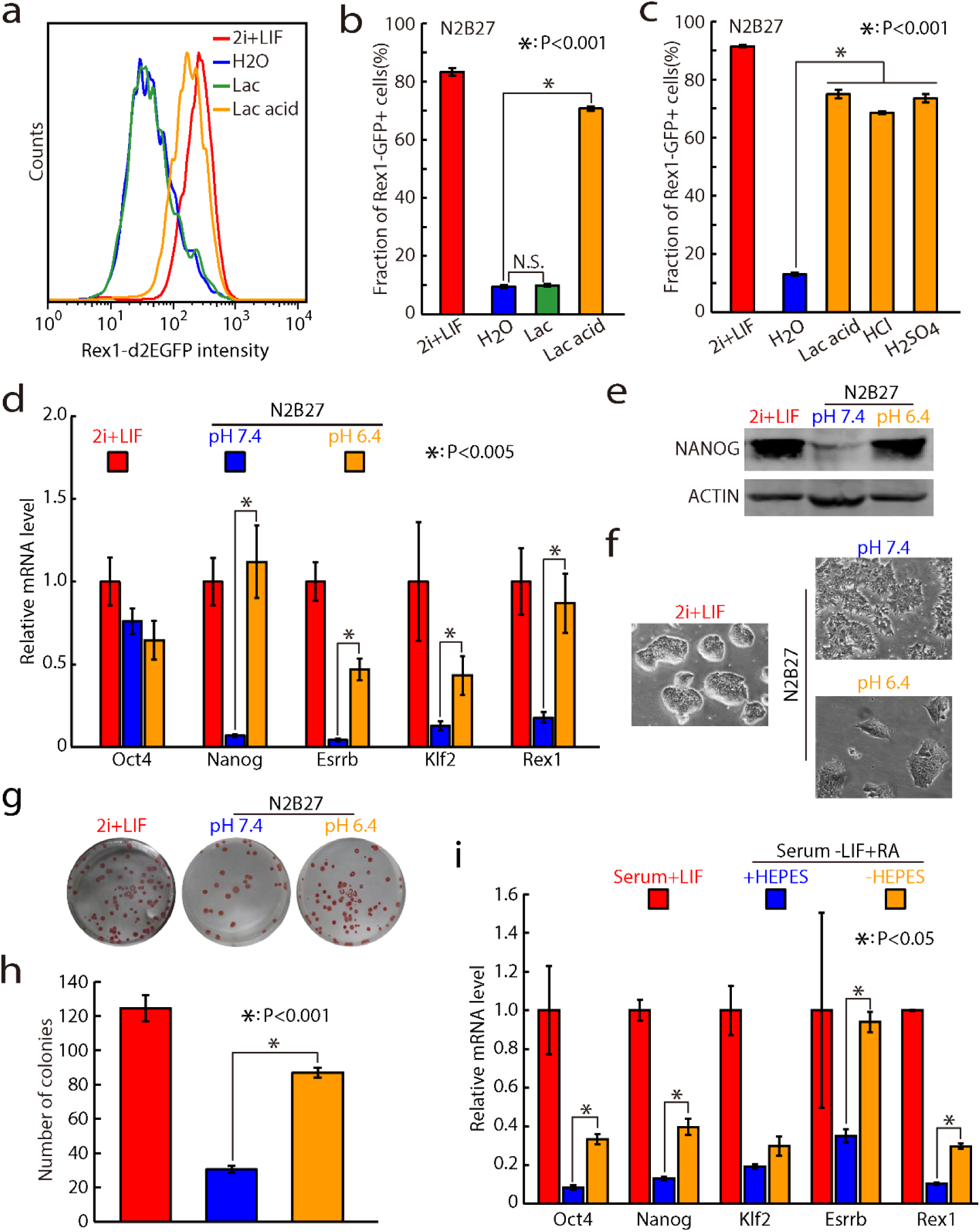
Lactic acid but not lactate promotes the self-renewal and pluripotency of mouse ESCs. (**a-c**) FACS analysis of GFP positive portions of Rex1-d2EGFP ESCs under different culture conditions; 10 mM lactic acid or lactate was used. (**d)** qRT-PCR analysis of pluripotency markers in low pH treated mouse ESCs; The Rpl7 gene was used as a control. Data were normalized to the mRNA level of 2i+LIF cultured mouse ESCs. Shown are mean ± SD, n = 3. (**e)** Western analysis of NANOG protein levels. (**f)** Morphology of low-pH treated mESC colonies. (**g, h)** Colony formation analysis. Representative images for whole well are shown for alkaline phosphatase (AP) staining. Colony numbers are shown as mean ± SD, n = 3. (**i**) qRT-PCR analysis of pluripotency markers in ESCs cultured in serum media with 1 μM retinoid acid and with or without HEPES buffer; The Rpl7 gene was used as a control. Data were normalized to the mRNA level of 2i+LIF cultured mouse ESCs. Shown are mean ± SD, n = 3.

To test what pH range provides the best effect, we grew Oct4-GFP-ires-Puro ESCs in differentiation media under various pH conditions and then evaluated for the maintenance of pluripotency by their ability to proliferate in 2i culture plus puromycin resistance^26^. The results show that acidic but not basic conditions promoted the self-renewal and pluripotency of ESCs with pH 6.0-6.4 being the most efficient (**Supplementary Fig. 1a**). Culturing ESCs at pH 5.5 or lower led to extensive cell death, making it impossible to evaluate their effects on pluripotency. We then further verified that pH 6.4 blocked the exit of pluripotency by flow cytometry analysis of Rex1-d2EGFP levels (**Supplementary Fig. 1b, c**) and qRT-PCR analysis of pluripotency and differentiation markers (**Fig. 1d**). Consistent with these results, NANOG protein was also highly expressed in N2B27-pH6.4 condition (**Fig. 1e**). Furthermore, ESCs cultured in N2B27-pH 6.4 retained a compact colony morphology, which indicates an undifferentiated state (**Fig. 1f**). Likewise, colony formation assay also confirmed the blocking of differentiation by pH 6.4 (**Fig. 1g, h**). To check the generality of acidic media blocking differentiation, we performed similar experiments in conventional serum media. Consistently, RT-PCR analysis showed that acidic condition blocked the differentiation of ESCs in serum media without LIF supplement as well (**Supplementary Fig. 1d**). Together, these results show that acidic microenvironment due to the accumulation of lactic acid promotes the self-renewal and pluripotency of mouse ESCs.

To test whether endogenously produced lactate is sufficient to support the self-renewal and pluripotency of mouse ESCs, we grew ESCs in serum media with 1 μM retinoid acid (RA) with or without HEPES buffer. After growing for approximately 48 hours, the pH of unbuffered media is decreased from 7.4 to 6.3 (**Supplementary Fig. 1e**), suggesting endogenously produced lactate sufficient to change the pH of extracellular environment. In contrast, the pH of buffered media remained constantly at 7.4. Consistent with pH changes, qPCR assay showed that ESCs cultured in unbuffered media retained high expression of pluripotency markers while ESCs in buffered media are differentiated (**Figure 1i**). Altogether, these data show that glycolysis can produce sufficient amount of lactate to block ESC differentiation.

### Transcriptomic and epigenomic analyses confirm that acidic condition maintains the pluripotency of ESCs

To fully characterize ESCs cultured in acidic media, we performed RNA-Seq, small RNA-Seq and ChIP-Seq using antibodies against OCT4, H3K27 tri-methylation (H3K27me3) and H3K4 tri-methylation (H3K4me3) for ESCs cultured in 2i+LIF-pH 7.4, 2i+LIF-pH 6.4, N2B27-pH 7.4 and N2B27-pH6.4. Principle component analysis (PCA) and unsupervised hierarchical clustering analysis of transcriptome (**Fig. 2a, Supplementary Fig. 2a, b**) and miRNA profiles (**Fig. 2b, Supplementary Fig. 2c**) showed that N2B27-pH 6.4 cultured ESCs clustered closer to pH 7.4 and pH 6.4 2i+LIF ESCs and are evidently separated from differentiated ESCs in N2B27-pH 7.4. As expected, low pH significantly inhibited the expression of differentiation driving MEK and GSK3 pathway genes and promoted the expression of pluripotency supporting AKT and STAT3 pathway genes (**Supplementary Fig. 2d**). These data support that acidic pH maintains the self-renewal and pluripotency of ESCs.

**Figure 2.**
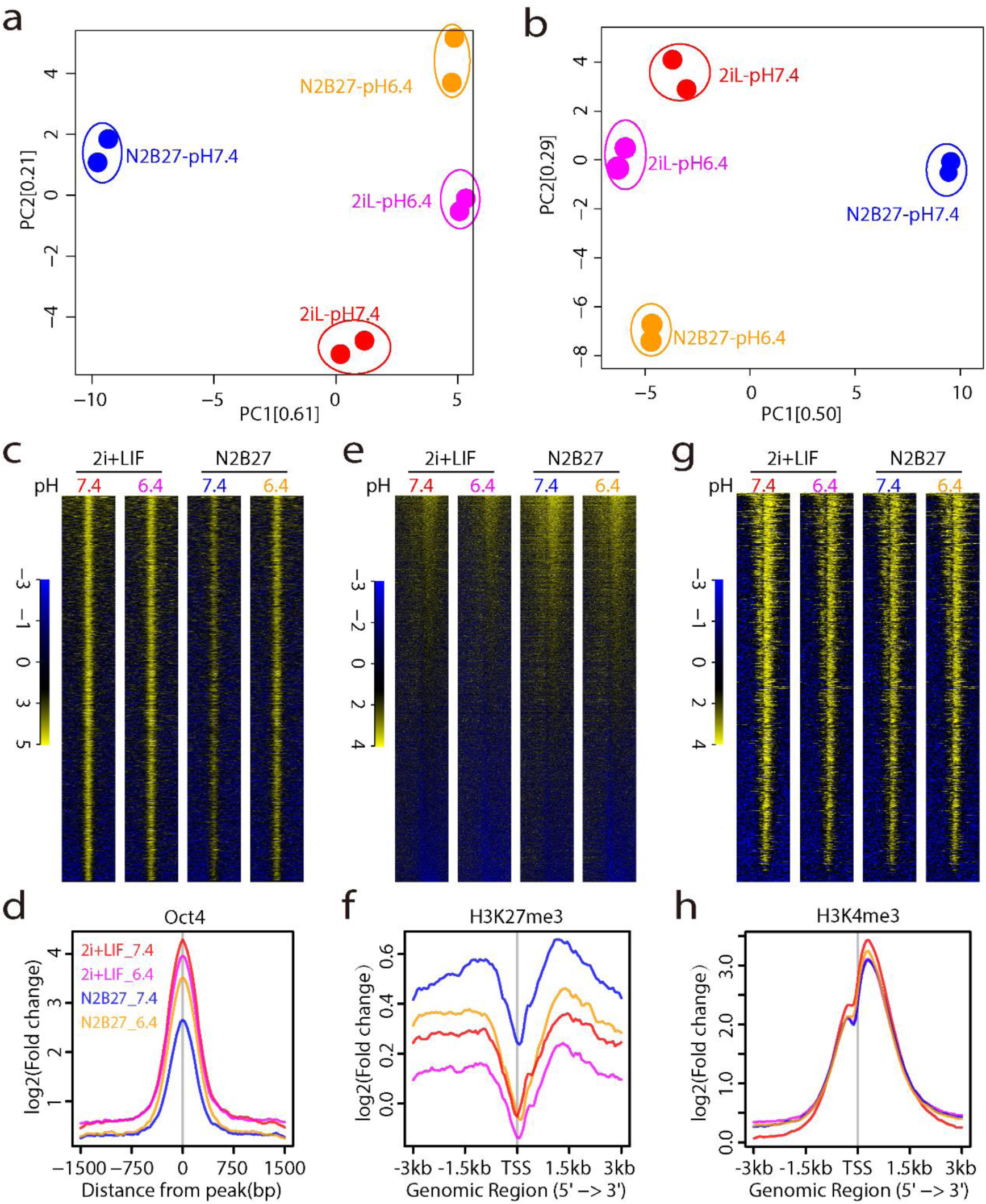
Transcriptomic and epigenomic analyses confirm that acidic condition maintains the pluripotency of ESCs. (**a**) PCA analysis for selected genes that are related to pluripotency and differentiation. (**b**) PCA analysis of global miRNA expression. (**c, d**) Heatmap and signal intensity plot of OCT4 ChIP. Shown are ±1.5 kb regions around the peak center of all peaks called by MACS2 (p value < 0.01); (**e, f**) Heatmap and signal intensity plot of H3K27me3 ChIP. Shown are promoters containing H3K27me3 peaks called by MACS2 (p value < 0.01) over random distribution in at least one of the culture conditions. (**g, h**) Heatmap and signal intensity plot of H3K4me3 ChIP. Shown are promoters containing H3K4me3 peaks called by MACS2 (p value < 0.01) over random distribution in at least one of the culture conditions.

Oct4 is one of the most important pluripotency regulators sitting at the center of the pluripotency transcriptional network^27^. We next asked whether Oct4 genomic occupancy was altered in N2B27-pH 6.4 ESCs. ChIP-Seq analyses showed that OCT4 binding remained high in N2B27-pH6.4 but is significantly downregulated in N2B27-pH 7.4 at global level (**Fig. 2c, d**). Importantly, ChIP-qPCR and ChIP-seq analyses showed that low pH rescued OCT4 binding at the promoter of many important pluripotency genes (**Supplementary Fig. 2e, f**). Furthermore, ChIP-seq analysis of H3K27me3 showed a global upregulation of H3K27me3 at transcriptional start sites in ESCs switched from 2i+LIF-pH7.4 to N2B27-pH 7.4 (**Fig. 2e, f**), consistent with substantial reorganization of H3K27me3 markers during ESC differentiation^28^. As expected from cellular phenotypes and transcriptomic analyses, this trend was evidently blocked by low pH treatment (**Fig. 2e, f and Supplementary Fig. 2g**). In contrast, we did not observe any significant differences in H3K4me3 profiles between all four conditions (**Fig. 2g, h**). Altogether, these data show that low pH treated mESCs are essentially similar to naive mESCs cultured in 2i+LIF at both transcriptomic and epigenomic levels.

### Low pH plus LIF maintain the long-term self-renewal of mouse ESCs

We then tested whether basal media at pH 6.4 can provide an alternative culture media to maintain and expand ESCs for long-term. Unfortunately, ESCs cultured in N2B27 at pH 6.4 could only be passaged for 2-3 passages and then quickly become degenerated (data not shown). Adding LIF to media helped to maintain ESC cultures for a few more passages. In addition, ESCs can be cultured in basal media at pH 6.8 with LIF for approximately one month, which were then gradually degenerated. These ESCs retained features of pluripotent ESCs, including compact colony morphologies (**Supplementary Fig. 3a**) and high expression of pluripotency markers (**Supplementary Fig. 3b**), while ESCs cultured in basal media at pH 7.4 plus LIF were mostly differentiated. These data demonstrate that basal media at pH 6.8 plus LIF can maintain self-renewal and pluripotency of mESCs for a relatively long period.

### Low pH promotes the self-renewal and pluripotency of human ESCs

Next, we tested whether low pH also promotes the self-renewal and pluripotency of human ESCs. Like mouse ESCs, human ESCs also heavily depend on aerobic glycolysis for energy supply^9, 13^. Strikingly, pH 6.4 media without bFGF partially maintained the self-renewal of hESCs as gauged by cell morphology (**Fig. 3a**), and the expression of core pluripotency genes OCT4, SOX2 and NANOG (**Fig. 3b**). In addition, flow cytometry analysis of OCT4-GFP human ESCs showed that low pH maintained high activity of OCT4 promoter (**Fig. 3c, d**). Together, these results indicate that the pluripotency promoting function of acidic microenvironment is conserved for both mouse and human ESCs.

**Figure 3.**
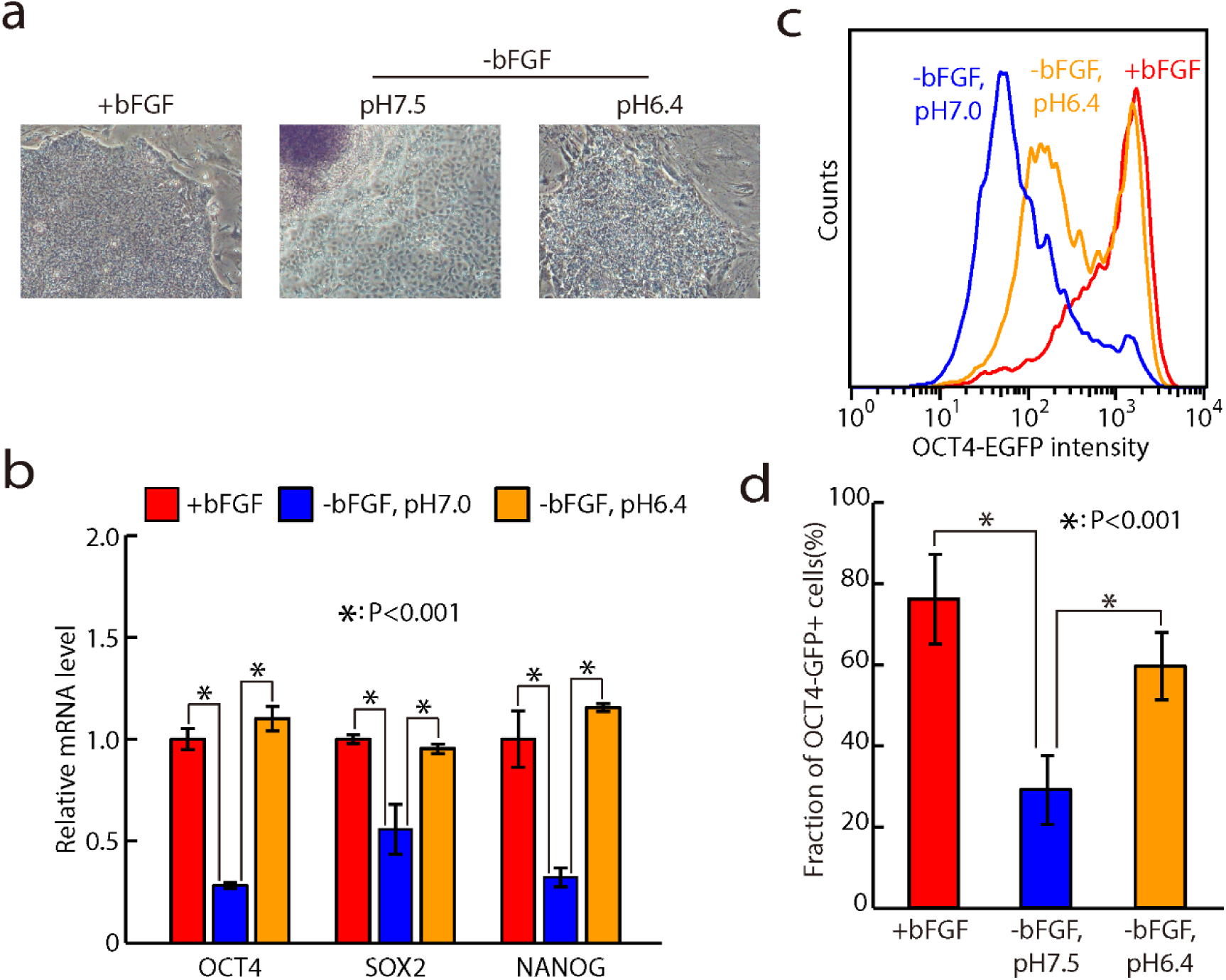
Low pH promotes the self-renewal and pluripotency of human ESCs. (**a**) Morphology of low-pH treated human ESCs. (**b**) qRT-PCR analysis of pluripotency markers of low-pH treated human ESCs. The β-actin gene was used as a control. Data were normalized to the mRNA level of bFGF cultured human ESCs. Shown are mean ± SD, n = 3. (**c, d**) FACS analysis of GFP positive portions of OCT4-GFP reporter human ESCs under different culture conditions.

### Low pH cause global reduction of H3K27 trimethylation at transcription start sites

Next, we sought to understand molecular mechanisms by acidic pH to promote the self-renewal and pluripotency of ESCs. As shown in **Figures 2c** and **2d**, we observed significant reduction of H3K27 tri-methylation at transcriptional start sites at low pH conditions. To assess the functional impact of H3K27me3 reduction, we analyzed the expression of H3K27me3 target genes and found they are significantly upregulated compared to all genes (**Fig. 4a**). In addition, for promoters that gained more H3K27me3 signals during differentiation, overall H3K27me3 intensity was significantly reduced in low pH cultures (**Fig. 4b, c**). Based on these data, we reasoned that if the reduction of H3K27me3 is responsible for the blocking of differentiation at low pH conditions, inactivating *Eed*, a gene essential for the tri-methylation of H3K27, should also block the differentiation of ESCs. To test this hypothesis, we generated *Eed* knockout ESCs using CRISPR/Cas9 (**Supplementary Fig. 4a-c**). Interestingly, genes upregulated in *Eed* knockout ESCs were significantly enriched in low pH treated ESCs (**Fig. 4d, e**), suggesting that low pH upregulates gene expression at least partially through the reduction of the tri-methylation of H3K27. Importantly, compared to wild type ESCs, *Eed* knockout ESCs displayed higher expression of pluripotency genes in N2B27 differentiation conditions (**Fig. 4f and Supplementary Fig. 4d**). Nevertheless, the effect of blocking differentiation by knocking out *Eed* was not as significant as by low pH treatment. Taken together, these data show that acidic environment maintains the pluripotency and self-renewal of ESCs at least partially by reducing the tri-methylation of H3K27 at transcriptional start sites.

**Figure 4.**
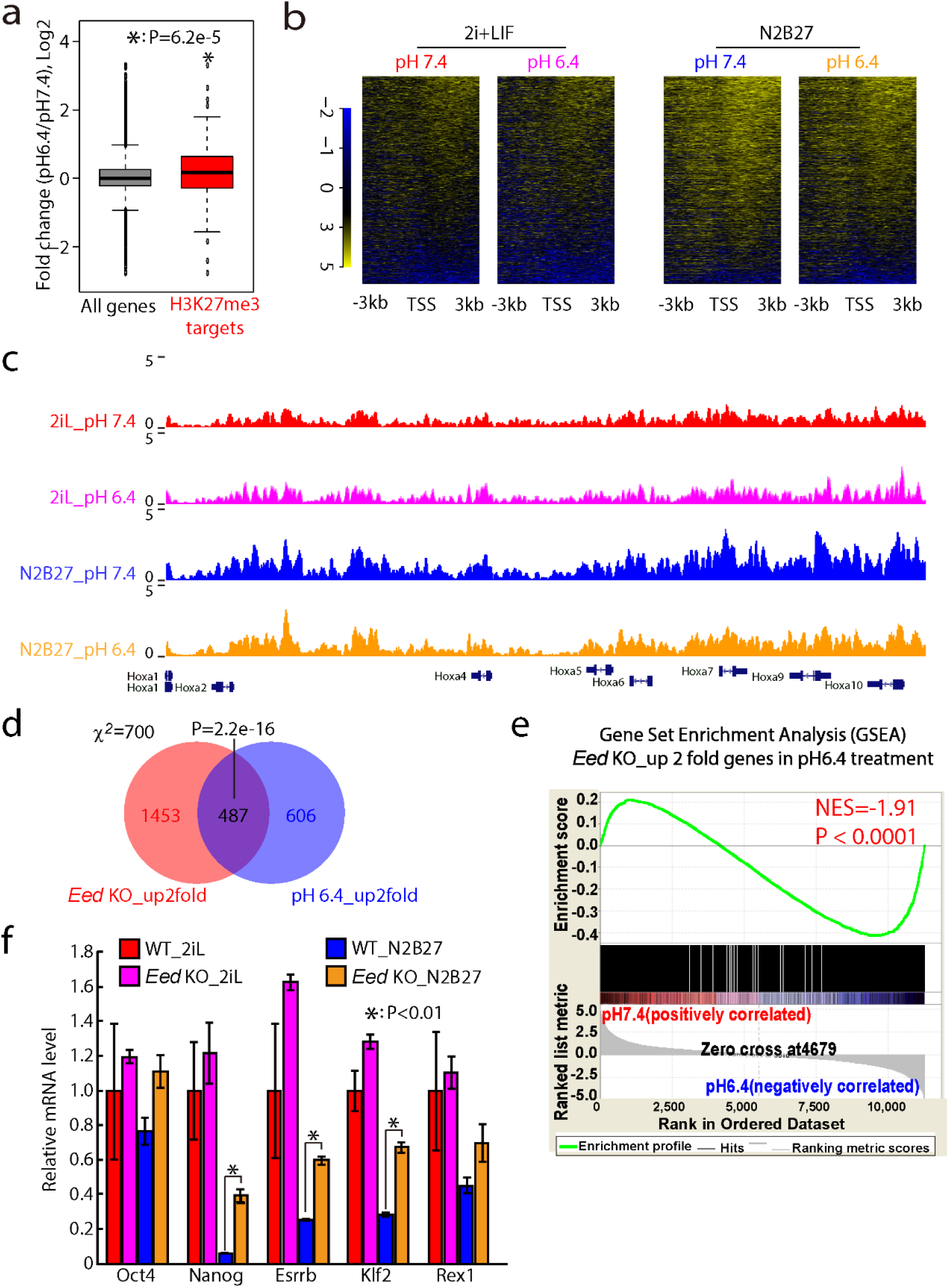
Low pH causes global reduction of H3K27me3 at transcriptional start sites. (**a**) Box plot for log2 fold change of mRNA expression for all expressed genes and genes that have H3K27 me3 peaks in 2i+LIF cultured mESCs. (**b**) Heatmap of H3K27me3 ChIP signals for promoters gained H3K27me3 signals during differentiation. (**c**) Genome browser screenshot of the H3K27me3 ChIP reads aligning to Hoxa locus. (**d**) Venn diagrams showing the overlaps between genes upregulated in *Eed* knockout mESCs and in pH6.4 treated mESCs. (**e**) GSEA analysis for >2 fold upregulated genes in *Eed* knockout mouse ESCs between pH7.4 and pH6.4 2i+LIF culture conditions. (**f**) qRT-PCR analysis of pluripotency markers in wild-type mESCs and *Eed* KO mouse ESCs differentiated for 1 day in N2B27 media without 2i and LIF. The β-actin gene was used as a control. Data were normalized to the mRNA level of 2i+LIF cultured mESCs. Shown are mean ± SD, n = 2.

### Low pH stabilizes thousands of mRNAs including known pluripotency genes in ESCs

Since knocking out *Eed* did not fully recapitulate the effects of low pH, we decided to search for other mechanisms by low pH in promoting the self-renewal and pluripotency of ESCs. Recent studies suggest that post-transcriptional regulation especially mRNA degradation is essential in controlling the pluripotency and differentiation of ESCs^29–32^. To check whether there is a link between low pH treatment and post-transcriptional regulation, we analyzed mRNA stability in low pH and control media. These experiments were performed in 2i+LIF media in order to exclude any secondary effects due to differentiation. The mRNA life time profiling was performed by collecting and analyzing RNA-seq data obtained at 0, 2, 4 and 8 hours after adding actinomycin D, a potent inhibitor of RNA polymerase II (**Fig. 5a**). Interestingly, accumulation plot showed that mRNAs were globally stabilized in low pH treated cells **Fig. 5b**). mRNA half-lives of 605 and 187 genes were increased or decreased by two fold (**Fig. 5c**), respectively. qRT-PCR analysis of randomly selected genes confirmed mRNA stabilization effects of low pH treatment (**Fig. 5d and Supplementary Fig. 5a**). As expected, overall expression level of stabilized genes and destabilized genes are upregulated and downregulated (**Supplementary Fig. 5b, c**), respectively.

**Figure 5.**
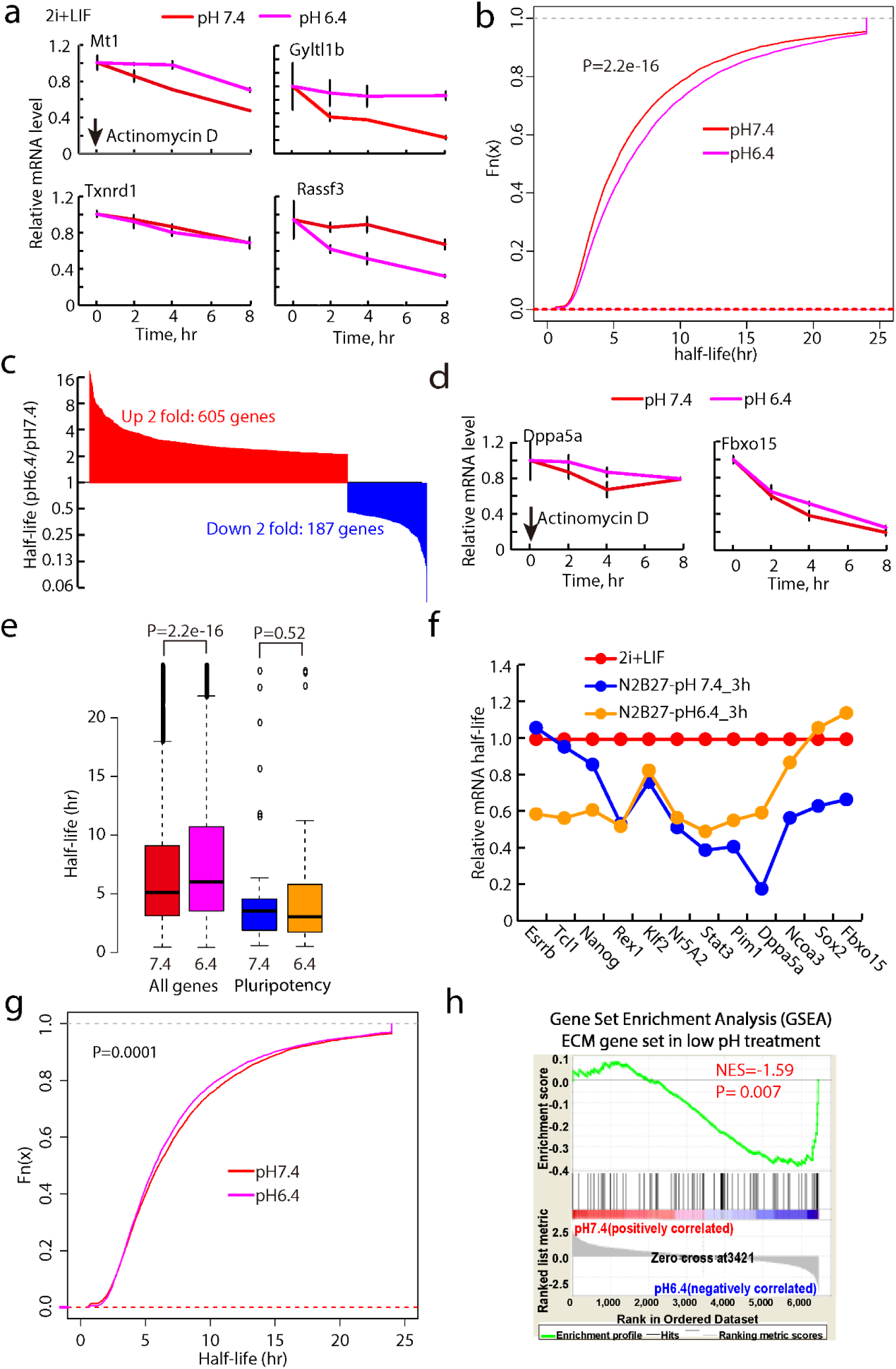
Low pH stabilizes thousands of mRNAs including known pluripotency genes. (**a**) mRNA degradation plot of representative genes. The β-actin gene was used as a control. For each gene, data were normalized to ESCs cultured in each condition without actinomycin D treatment (0 hour). Shown are mean ± SD, n = 2. Actinomycin D was added 48 hr after low pH treatment, and cells were collected at 0, 2, 4, and 8 hr after actinomycin D addition for RNA-seq. (**b**) Cumulative plot for mRNA half-life of ESCs in pH7.4 and pH6.4 culture conditions. (**c**) Bar plots of log2 fold change of mRNA half-life between pH6.4 and pH7.4. (**d**) Half-life of represented pluripotency genes Dppa5a and Fbxo15. The β-actin gene was used as a control. For each gene, data were normalized to ESCs cultured in each condition without actinomycin D treatment (0 hour). Shown are mean ± SD, n = 2. (**e**) Box plot for mRNA half-life of pluripotency genes in pH7.4 and pH6.4 culture conditions. (**f**) mRNA half-life analysis of pluripotency genes in ESCs after cultured in N2B27 media for 3 hours at pH 7.4 and pH 6.4. Data were normalized to mRNA half-life of each gene in 2i+LIF ESCs. (**g**) Cumulative plot for mRNA half-life of U2OS cells in pH7.4 and pH6.4 culture conditions. (**h**) GSEA analysis for ECM gene set of U2OS cells in pH7.4 and pH6.4 culture conditions.

We then analyzed whether mRNAs of pluripotency genes are stabilized. Dppa5a and Fbxo15 were slightly stabilized in 2i+LIF pH 6.4 versus 2i+LIF pH 7.4 ESCs (**Fig. 5d**). However, overall mRNA half-lives for pluripotency genes were similar in 2i+LIF cells under normal or low-pH conditions (**Fig. 5e**). These results suggest: 1. low-pH has no effects in stabilizing pluripotency genes; or 2. low pH has effects in stabilizing pluripotency genes, but other mechanisms stabilizing pluripotency genes in 2i+LIF condition cover up the function of low-pH treatment. If the latter is right, mRNA of pluripotency genes should become destabilized upon removing 2i+LIF and low pH treatment should block this tendency. To test this, we analyzed mRNA half-life three hours after removing 2i and LIF, at which point N2B27-pH 7.4 cultured ESCs are still pluripotent. Interestingly, mRNA half-lives of 11 out of 12 selected pluripotency genes were significantly decreased, and low-pH stabilized 6 of them including Stat3, Pim1, Dppa5a, Ncoa3, Sox2 and Fbxo15 (**Fig. 5f**). Among them, overexpression of Stat3, Pim1 and Ncoa3 have been shown to block ESC differentiation in various cases^33–35^. These data suggest that low pH help maintain pluripotency state by stabilizing a subset of pluripotency mRNAs.

### Low pH stabilizes extracellular matrix genes in cancer cells

To test whether stabilization of mRNAs is a general mechanism by low pH treatment in other cells, we measured mRNA half-life in U2OS cells cultured under normal and low pH media. Consistent with previous findings in cancer cells^23, 36, 37^, low-pH treatment promoted epithelial-mesenchymal transition (EMT) in U2OS and HepG2 cells (**Supplementary Fig. 6a, b**). In these cells, we did not observe similar global stabilization of transcripts as in ESCs (**Fig. 5g**), instead we found around 300 genes are stabilized or destabilized (> 2 fold changes of half life) upon low pH treatment. However, we found that mRNAs of extracellular matrix (ECM) proteins are significantly stabilized upon low-pH treatment (**Fig. 5h and Supplementary Fig. 6c**). This is consistent with EMT observed in these cells upon low pH treatment since upregulation of ECM components are associated with EMT in cancer cells. Therefore, low-pH treatment could stabilize mRNAs of a subset of key genes to promote EMT. Together, these data suggest that mRNA stabilization might be a general mechanism by low pH environment to confer functional consequences in ESCs and cancer cells.

### Low pH downregulates AGO1 protein expression and derepresses a subset of miRNA targets in ESCs

miRNA is one of the most important post-transcriptional regulators that destabilizes mRNAs. To gain more insights on how low pH promotes ESC self-renewal and pluripotency, we considered the possibility that a subset of mRNAs are stabilized through the inhibition of miRNA activities. We focused on miR-294/302 family of miRNAs, which are the most enriched miRNAs in ESCs. We first defined their targets based on their 3’UTR sequences and their expression changes in *Dgcr8* KO versus wild type ESCs (see **Materials and Methods**). Interestingly, we found mRNA targets of miR-294/302 family were significantly enriched in stabilized genes (**Fig. 6a and Supplementary Fig. 7a**). Consistently, GSEA showed that miR-294/302 targets are significantly upregulated in low pH treated ESCs (**Fig. 6b and Supplementary Fig. 7b**). RT-PCR confirmed the upregulation of previously verified miR-294 targets Fndc3b, Mbd2 and Tgfbr2 and the downregulation of Mbd2 target Myc (**Fig. 6c**). In addition, luciferase reporter assay showed that the inhibition at the 3’UTR of Mbd2 is partially reversed by low pH treatment (**Fig. 6d**). In the next, we verified that inhibition of miRNA activities indeed can significantly block ESC differentiation in the absence of 2i and LIF using *Dgcr8* knockout ESC (**Fig. 6e**). These results are also consistent with our previous reports showing that miR-294/302 family is important for the efficient exit of naive pluripotency and subsequent differentiation of ESCs^38, 39^. Therefore, the de-repression of miRNA targets can partially explain the blocking of differentiation by low pH condition

**Figure 6.**
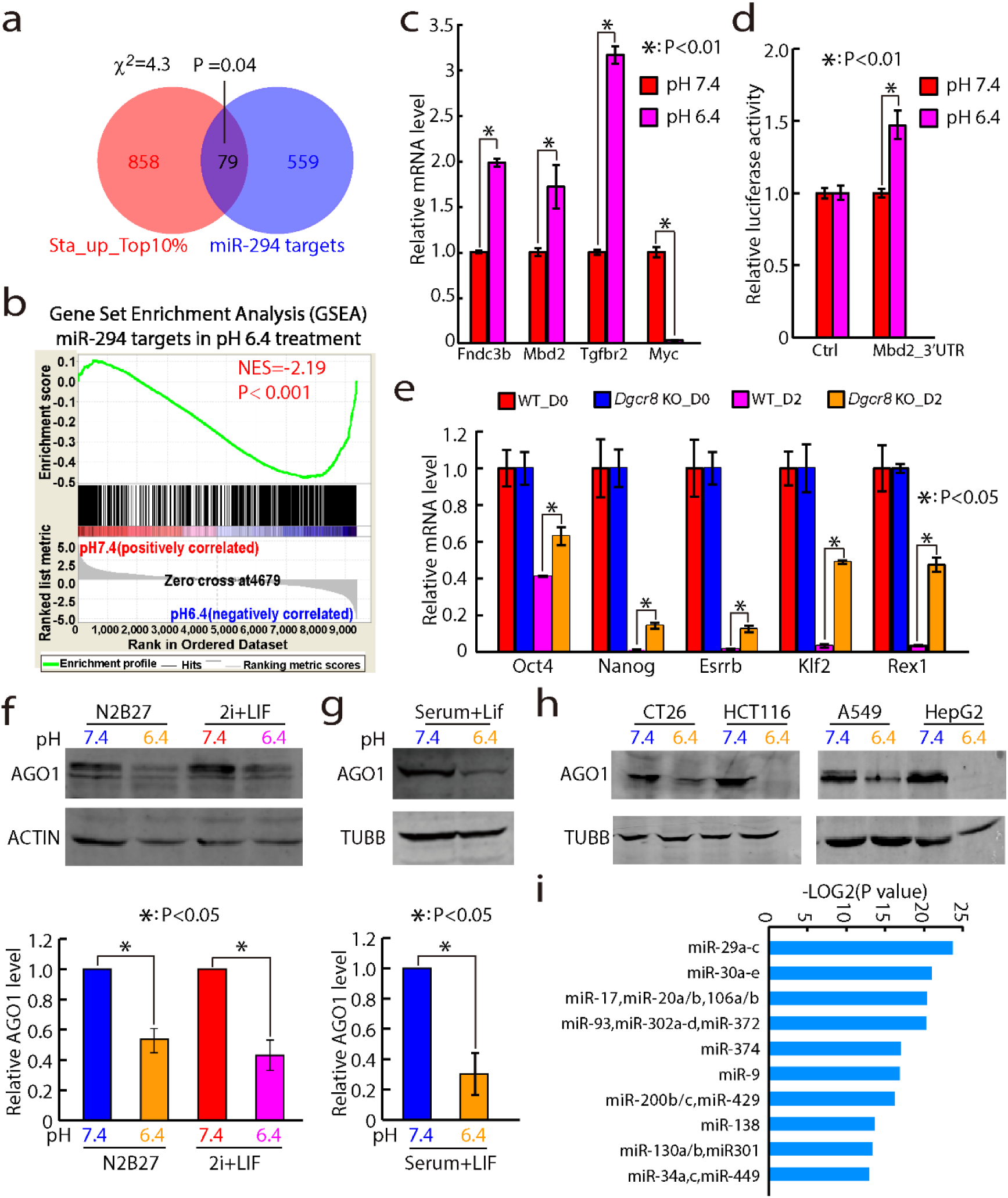
Low pH downregulates AGO1 protein expression and derepresses a subset of miRNA targets. (**a**) Venn diagrams showing the overlaps between genes with extended mRNA half-lives in pH6.4 condition and miR-294 target genes. (**b**) GSEA analysis for miR-294 target genes in pH7.4 and pH6.4 culture conditions. (**c**) qRT-PCR analysis of canonical miR-294 target genes (Fndc3b, Mbd2 and Tgfbr2) and Myc expression in low-pH treated mESCs; The Rpl7 gene was used as a control. Data were normalized to the mRNA level of 2i+LIF cultured mouse ESCs. Shown are mean ± SD, n = 3. (**d**) Luciferase reporter assay. Shown are mean± SD, n = 3. (**e**) qRT-PCR analysis of pluripotency markers in wild-type ESCs and *Dgcr8* knockout mouse ESCs cultured for 2 days in N2B27 differentiation media. The β-actin gene was used as a control. Data were normalized to the mRNA level of 2i+LIF cultured mouse ESCs. Shown are mean ± SD, n = 2; (**f, g**) Western analysis AGO1 in mESCs cultured in different pH conditions. The ACTIN or TUBBLIN was used as a loading control. (**h**) Western analysis AGO1 in human cancer cell lines U2OS and HepG2 in different pH condition. The TUBBLIN was used as a loading control. (**i**) GSEA of miRNA targets for genes upregulated (2 fold) in low pH treated U2OS cancer cells.

The de-repression of miRNA targets could be due to the decrease of miRNA expression or the decrease of the executer of miRNA repression (i.e. Argonaut proteins). The miR-294/302 family includes 6 members (miR-291a/294/295-3p and miR-302a/b/d-3p) with the same seed sequence. miRNA profiling data and qRT-PCR showed that the total level of miR-294/302 family was not significantly altered at low pH conditions (**Supplementary Fig. 7c, d**). We then checked the mRNA and protein level of AGO1 and AGO2, since AGO3 and AGO4 were not expressed in ESCs. Interestingly, while both protein and mRNA of Ago2 changed little, AGO1 protein but not Ago1 mRNA was significantly downregulated in 2i+LIF ESCs upon low pH treatment (**Fig. 6f and Supplementary Fig. 7e, f**). We further confirmed that AGO1 protein was also downregulated in conventional serum cultured ESCs (**Fig. 6g**). More interestingly, low pH treatment also led to significant reduction of AGO1 protein in various human cancer cell lines (**Fig. 6h**). The reduction of AGO1 is consistent with EMT promoting effects of low pH, since miRNAs including miR-200, miR-29 and miR-34 families have been shown to suppress EMT in various cancer^40^. Indeed, we found that mRNA targets of these miRNAs are enriched in upregulated genes in low pH treated U2OS cells (**Fig. 6i**). Altogether, these data show that low pH downregulates AGO1 protein to de-repress a subset of miRNA targets in ESCs and cancer cells.

## Discussion

Why PSCs prefer aerobic glycolysis over oxidative phosphorylation is not clear. While previous research has focused on identifying intermediate metabolites that are essential for the epigenetic regulation of pluripotency^1, 2, 14, 17^, the function of microenvironment shaped by lactic acidosis is largely overlooked. Here we present evidence that acidic microenvironment due to lactate production helps maintain the self-renewal and pluripotency of mouse and human ESCs (**Fig. 7**). RNA-Seq, miRNA Seq and OCT4, H3K4me3 and H3K27me3 ChIP-Seq confirmed that low-pH treated ESCs are indeed pluripotent at transcriptomic and epigenomic levels. Mechanistically, we showed that low-pH treatment maintains ESC self-renewal and pluripotency through both epigenetic and post-transcriptional mechanisms. First, the tri-methylation of H3K27 is globally reduced at acidic conditions. Second, acidic pH stabilized hundreds of mRNAs including many pluripotency genes. In addition, we found that AGO1 protein is downregulated at acidic pH, leading to the global upregulation of miR-290/302 targets in ESCs. Importantly, we found that AGO1 downregulation and mRNA stabilization by low-pH treatment are also conserved in cancer cells, indicating the conservation of a common pathway mediating low pH response in different cell lineages. Altogether, our study provides insights into mechanisms whereby acidic microenvironment shaped by enhanced glycolysis regulates gene expression and cell fate. In addition, our findings have broad implications for the fields of regenerative medicine and cancer.

**Figure 7.**
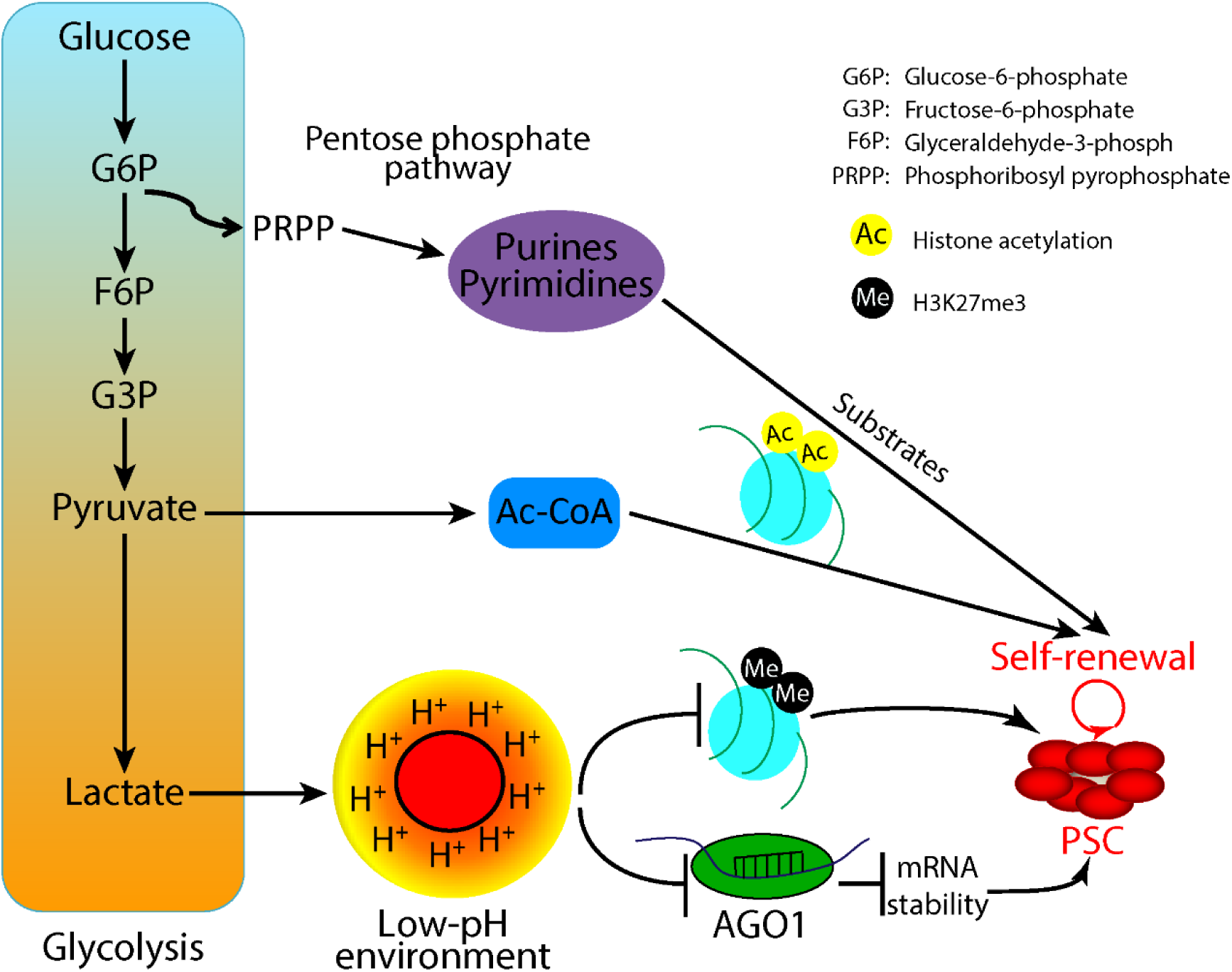
Summary graph showing that lactic acid generated from glycolysis promotes the self-renewal and pluripotency of ESCs.

The pH range 6.4-6.8 used in this study is physiologically relevant for mouse and human blastocysts as well as human tumors^20, 41^. Acidosis is associated with multiple malignant phenotypes in tumors including increased metastasis and radioresistance^21, 22^. In contrast, the role of acidic environment in blastocysts remains elusive. Our in vitro study points to an interesting hypothesis that lactic acid may work together with other signaling pathways to maintain the self-renewal and pluripotency of epiblast cells during early embryonic stages. Consistent with this, embryos passing 8-cell stage, approximately when the metabolism of an embryo shifts to glycolysis, prefers culture media with lower pH for full development^42^. In addition, *Ldha* knockout leads to preimplantation death in mouse embryos (The Sanger Mouse Genetics Program). Future work should elucidate the functional roles of lactate accumulation in embryo development. These studies may lead to the development of better means to culture embryos for in vitro fertilization (IVF) practice.

Enhanced glycolysis is observed in a variety of stem cells and cancer cells^1–5^. Elucidating its function and working mechanism can provide insights on how to manipulate stem cells and cure cancers. However, directly manipulating pH in vivo to change the microenvironment of stem cells and cancers is apparently not feasible. Therefore, understanding the downstream molecular impact of extracellular acidity is of high importance, as this will identify candidate pathways that may be manipulated to control stem cells or cancer growth. Our study for the first time makes connections between the extracellular acidity and post-transcriptional regulation in ESCs and cancer cells. RNA binding proteins, miRNAs as well as mRNA modifications (e.g. m6A) are major post-transcriptional regulators that have demonstrated functions in stem cells and cancer cells^43–45^. Future studies are warranted to elucidate the components of these pathways and other novel pathways that link extracellular acidity to post-transcriptional regulation. Studying these components may lead to the invention of innovative approaches in regenerative medicine and cancer therapy.

## Materials and Methods

### Cell culture and differentiation assay

Mouse ESCs were cultured on 0.1% gelatin-coated plates in 2i+LIF media which is consist of N2B27 supplemented with PD0325901 (1.0 μM), CHIR99021 (3.0 μM) and leukemia inhibitory factor (LIF, 1000 unit /ml) or serum media which is consist of DMEM supplemented with 15% FBS and LIF. For differentiation assay, ESCs were first plated in 2i+LIF for 20 hr, then washed once with 1xPBS and changed to N2B27 with indicated supplements, cultured for another 2 days before analysis with media changed every day; For colony formation assay, 200 cells were plated on a well of 12-well on 0.1% gelatin-coated plate in 2i+LIF media. Human ESCs were cultured on MEF in KSR medium knockout DMEM with 20% KSR supplemented with 100 mM 2-mercaptoethanol (2ME), MEM nonessential amino acids (NEAA), 2 mM L-glutamine and 5ng/ml bFGF (Leto Laboratories, Beijing). For differentiation assay, 300 single hESCs were plated on MEF in KSR medium with bFGF adding Y27632(10uM), then bFGF and Y27632 were withdrawn second day and changed medium every two days until the clone is formulated. Immediately before media change, pH of media was adjusted by lactic acid, hydrochloride acid and sulphuric acid and measured by pH meter at room temperature.

### RNA extraction and qRT-PCR

Total RNA was extracted following standard Trizol protocol (Invitrogen). RT-PCR was performed using Sybr Green mix (Vazyme) with ABI Step One Plus (Applied Biosystems).

### RNA-seq, Small RNA-seq and bioinformatics analysis

RNA and small RNA libraries from two independent biological replicates were generated using NEBNext small RNA library Prep Set for Illumina (E7420) and NEBNext Small RNA Library Prep Set (E7330S) according to the manufacturer’s instruction, respectively. NEBNext multiplex oligos for Illumina (E7335) were used. The final libraries were measured by Qubit fluorometric assay (Life Technologies). Libraries were sequenced by Illumina HiSeq 2500. Data was analyzed as previously described^39^. R 3.1.1 was used for generation of the heatmap and PCA plot. List of selected genes related to pluripotency and differentiation is as previously published^46^. GSEA2.4 was used to test for the enrichment of selected gene sets by java GSEA Desktop Application.

### ChIP-qPCR and ChIP-seq

Cells under different culture conditions were cross-linked with 1% formaldehyde for 15 min at room temperature followed by the addition of 125 mM glycine to inactivate formaldehyde. Chromatin extracts containing DNA fragments with an average size of 200–500 bps were immunoprecipitated using IgG or antibodies against H3K27me3 (Abclonal, A2363), H3K4 me3 (Abclonal, A2357) and OCT4 (Santa, sc-8628) at 4°C overnight. ChIP-Seq library was generated by NEBNext Ultra DNA Library Prep Kit (E7370). Libraries were submitted for sequencing on Illumina Hiseq 2000. Sequenced reads after trimmed were mapped to mouse mm10 whole genome using STAR v2.3.0. Peaks were called by MACS2 with default parameters (p value ⩽0.01 and --SPMR). Intensity plot and heatmap were generated by ngs.plot.r with aligned BAM files. Peak plot was generated by IGV software.

### mRNA half-life analysis

To measure mRNA stability, transcription was blocked by adding actinomycin D (Sigma, cat # A-9415) to the medium at the concentration of 10 mg/ml. Cells were harvested at 0, 2, 4 and 8 h after the addition of actinomycin D and processed for RNA-seq or qRT-PCR preparation and analysis. For RNA half-life calculation, genes with average FPKM lower than 0.2 in all samples were deemed as lowly expressed and were removed from further analysis. Filtered genes were normalized to β-actin. Then a linear regression of the form y = a - bt was performed on each gene through the four time points, where y is the log-transformed (base 10) read count, t is the time, b is the slope, a is the intercept, and d = b x ln(10) is the instantaneous decay rate. Half-life of each gene was calculated and projected as H = min (24, ln(2)/d) for positive d and h = 24 hr for negative d. Genes that taken for further analysis were determined by requiring correlation (R) of linear data was < −0.7 in all samples.

### miR-294/302 target analysis

A gene is defined as a miR-294/302 target in ESCs based on two criteria: First, it has at least one predicted miRNA binding sites complementary to the seed sequence “AAGUGC”; Second, its expression is significantly upregulated in *Dgcr8* knockout versus wild type ESCs (top 10% ranked by fold changes > 1.99 fold). Similarly, top 10% genes with increased mRNA half-lives (> 1.83 fold) were used to make Venn diagram in Figure 6.

### Western blot analysis

Antibodies against NANOG from Calbiochem (Darmstadt, Germany), ACTIN (AP0060) and TUBBLIN (AP0064) was from Bioworld Technology (Nanjing, China). Anti-rabbit and mouse secondary antibodies was from LI-COR and membranes were imaged using Oddssay.

### Statistical analysis

The data were presented as mean ± SD except where indicated otherwise. We performed two-tailed unpaired Student’s t-test to determine statistical significance except for analysis shown in the boxplot graph (two-tailed Wilcoxon signed-rank test), cumulative plot (Kolmogorov-Smirnov test) and Venn diagram (chi-square test). P value < 0.05 was considered as statistically significant.

## Acknowledgements

We would like to thank members of Wang laboratory for critical reading and discussion of the manuscript. This study was supported by The National Key Research and Development Program of China [2016YFA0100701] and the National Natural Science Foundation of China [31471222, 31622033 and 31521062] to YW.

## Author Contributions

WTG performed most of experiments with help from other authors. SHW performed experiments in Figure 3. XSZ and XWW performed experiments related *Eed* knockout. KLG and JH helped with bioinformatics analysis. All authors were involved in the interpretation of data. YW and WTG conceived the project. YW supervised the project. YW and WTG wrote the manuscript with help from other authors.

## Availability of Data and Materials

All data generated or analysed during this study are included in the manuscript and its supplementary information files. Original sequencing data used and/or analysed during the current study are available from the corresponding author on reasonable request.

## SUPPLEMENTARY FIGURE LEGENDS

**Figure S1, Related to Figure 1.**
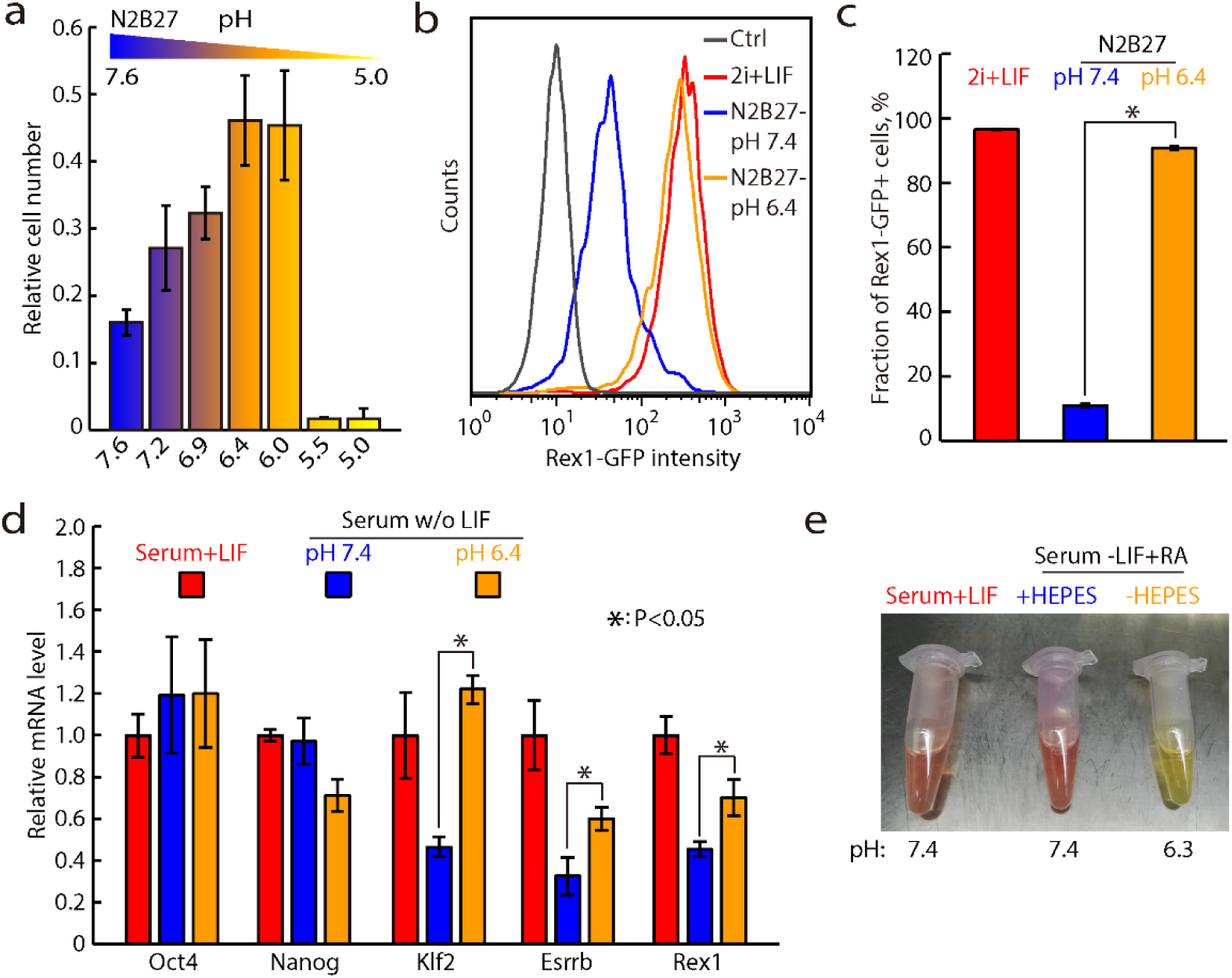
Low pH promotes the self-renewal and pluripotency of mouse ESCs. (**a**) MTT assay to test different pH conditions for maintaining pluripotency. OCT4-GFP-ires-Puro mESCs were exposed to N2B27 media at different pH for three days, and selected by 2 μg/ml puromycin in 2i media for two days. Shown are mean ± SD, n = 4. (**b, c**) FACS analysis of GFP positive portions of Rex1-GFP reporter cells in different culture conditions. (**d**) qRT-PCR analysis of pluripotency markers of low pH treated mESCs in Serum media; The Rpl7 gene was used as a control. Data were normalized to the mRNA level of Serum + LIF cultured mESCs. Shown are mean ± SD, n = 2. (**e**) pH value of media with or without HEPES buffer after growing ESCs for two days.

**Figure S2, Related to Figure 2.**
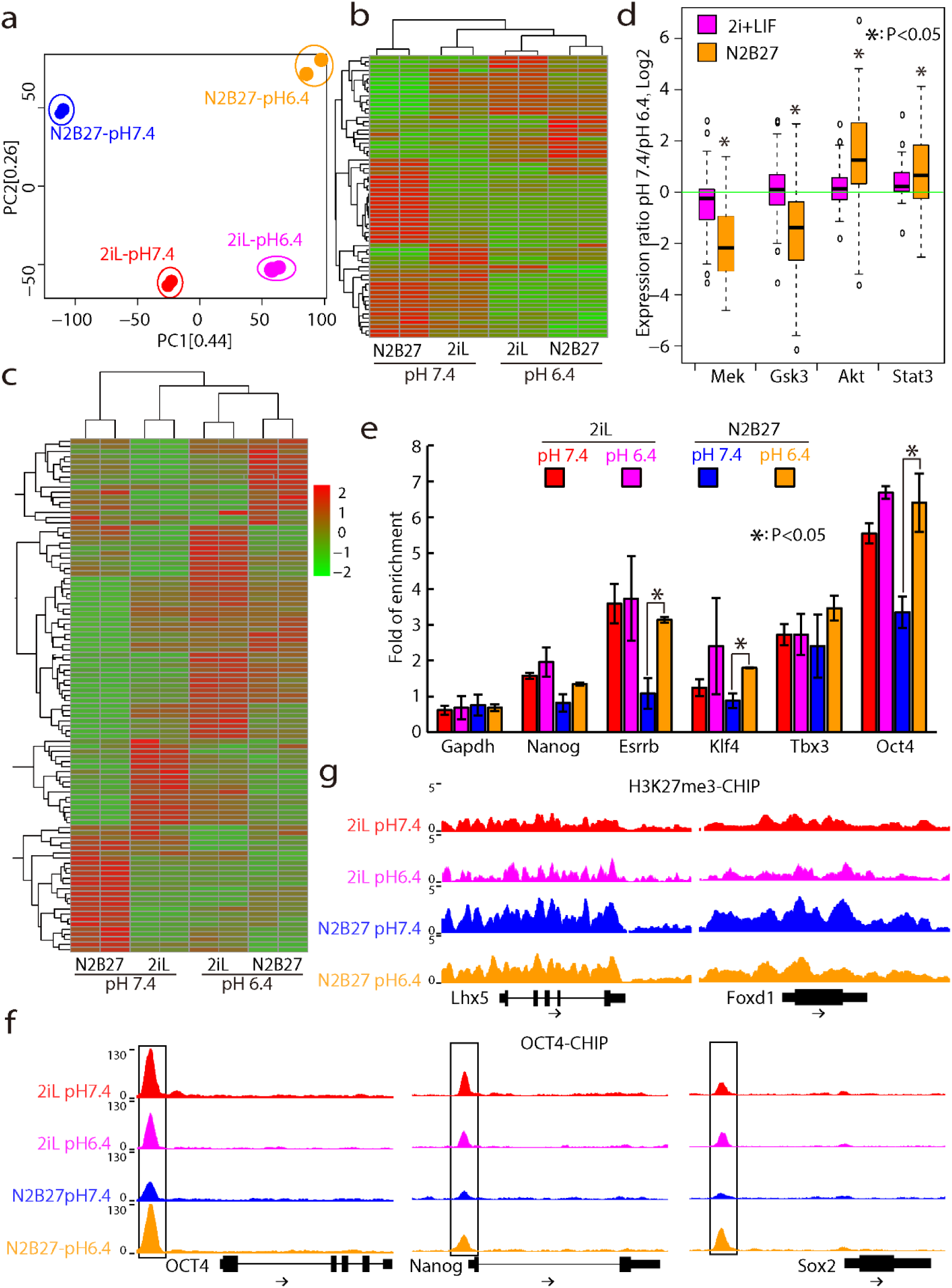
Low pH treated mESCs are similar to naive mESCs cultured in 2i+LIF at transcriptomic and epigenomic levels. (**a**) PCA analysis of global mRNA expression. (**b**) Heatmap of selected genes that are related to pluripotency and differentiation; (**c**) Heatmap of global miRNA expression. (**d**) Box plot showing pathway analysis for mESCs under normal pH and low-pH conditions. (**e**) ChIP-qPCR analysis of OCT4 binding at the promoter of pluripotency genes. (**f**) Screenshot of OCT4 binding of pluripotency genes *Oct4*, *Sox2* and *Nanog* under different culture conditions; (**g**) Screenshot of H3K27me3 ChIP reads mapping to the promoter of *Foxd1* and *Lhx5* under different culture conditions.

**Figure S3, Related to Figure 1 & 2.**
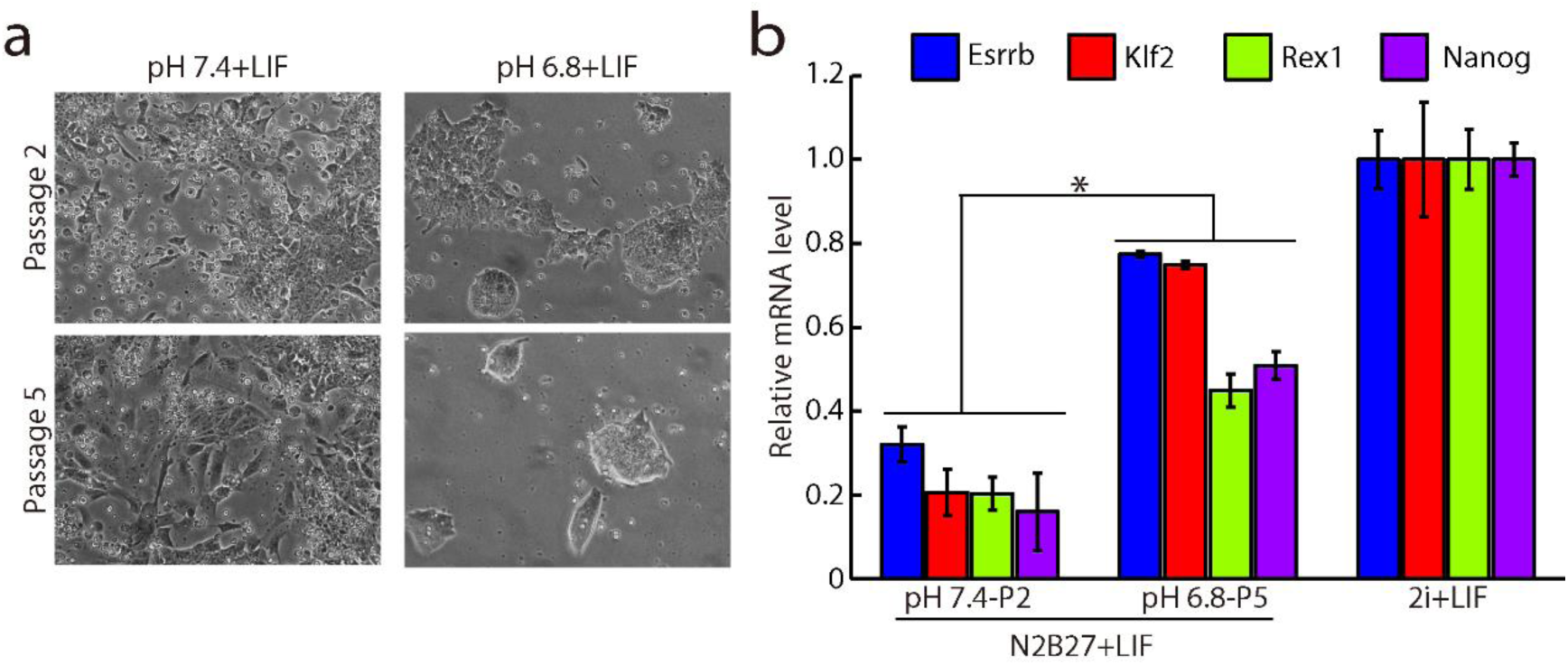
Low pH plus LIF supports long term self-renewal of mouse ESCs. (**a**) Morphology of passage 2 and 5 ESC colonies. (**b**) qRT-PCR of pluripotency genes for passage 2 LIF pH 7.4 cells and passage 5 LIF pH 6.8 cells. The Rpl7 gene was used as a control. Data were normalized to the mRNA level of 2i+LIF cultured mESCs. Shown are mean ± SD, n = 2.

**Figure S4, Related to Figure 4.**
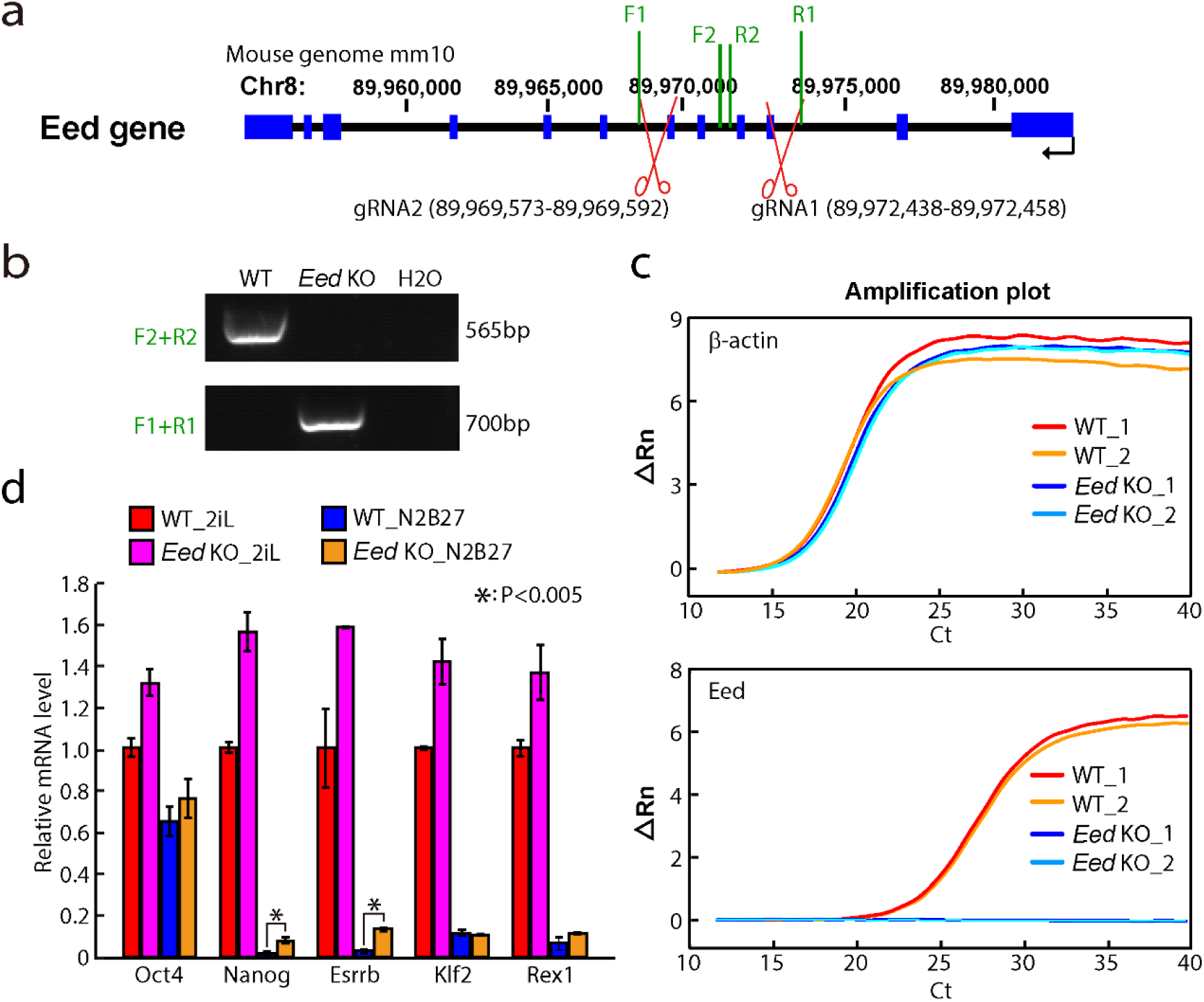
Generation of *Eed* knockout mouse ESCs. (**a**) Illustration of guide RNA (gRNA) sequences for knocking out Eed; location of primers used for genomic PCR confirming knockout of *Eed* are shown. (**b**) Genomic PCR of *Eed* knock out ESCs; (**c**) qRT-PCR amplification plot of β-actin and Eed in wild type and *Eed* knockout mouse ESCs. (**d**) qRT-PCR analysis of pluripotency markers in wild-type mESCs and *Eed* KO mouse ESCs differentiated for 2 days in N2B27 media without 2i and LIF. The β-actin gene was used as a control. Data were normalized to the mRNA level of 2i+LIF cultured mESCs. Shown are mean ± SD, n = 2.

**Figure S5, Related to Figure 5.**
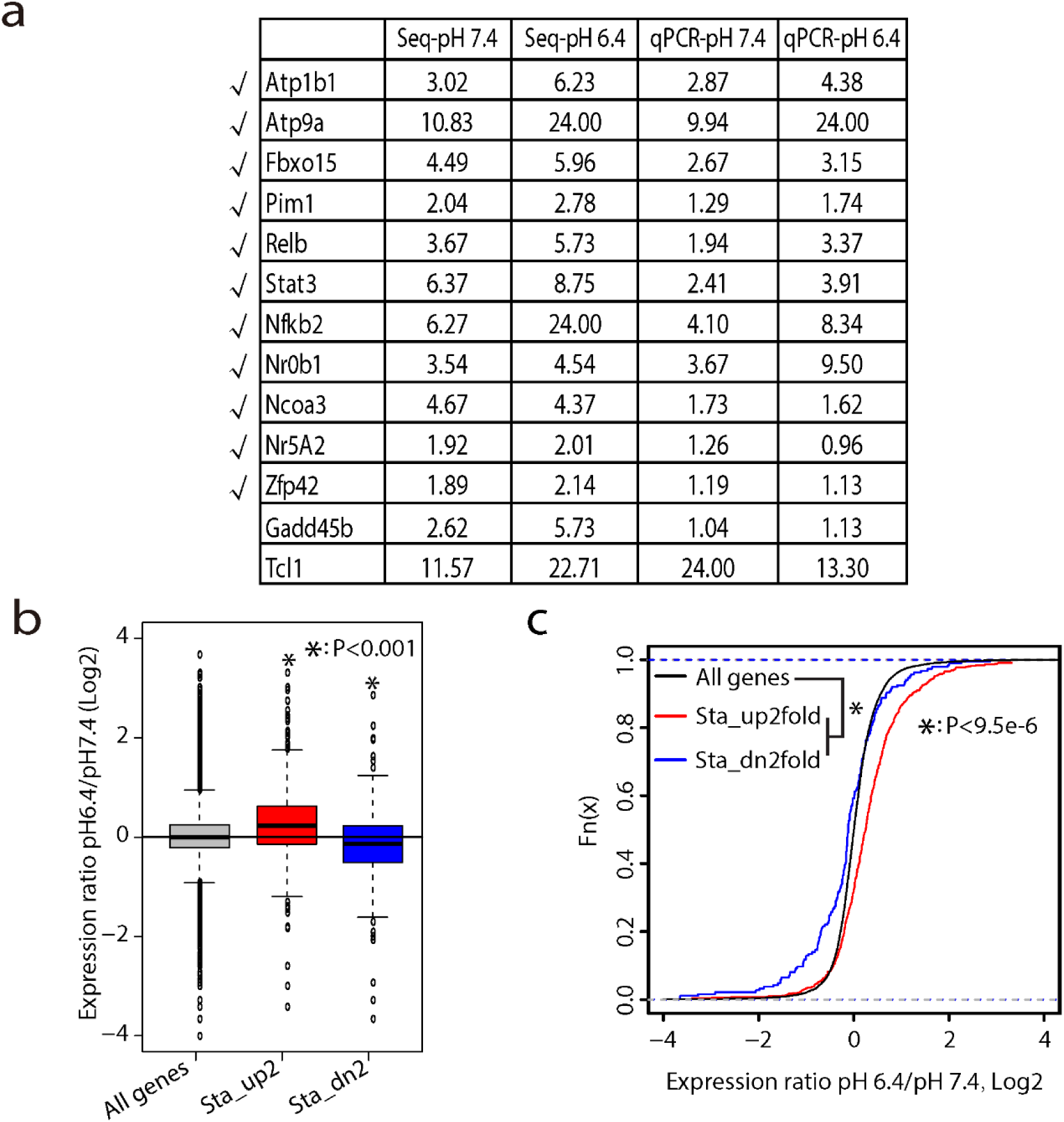
Acidic pH stabilizes a large number of genes including many pluripotency genes in ESCs. (**a**) mRNA half-life in RNA-seq data and qRT-PCR analysis for random picked genes; Genes with consist mRNA half-life between RNA-seq data and qRT-PCR analysis were labelled with “√”. (**b**) Box plot of Log2 fold changes (pH6.4/pH7.4) of mRNA expression for genes with half-life increased or decreased > 2 fold in pH6.4 media. (**c**) Cumulative plot of Log2 fold changes (pH6.4/pH7.4) of mRNA expression for genes with half-life increased or decreased > 2 fold in pH 6.4 media.

**Figure S6, Related to Figure 5.**
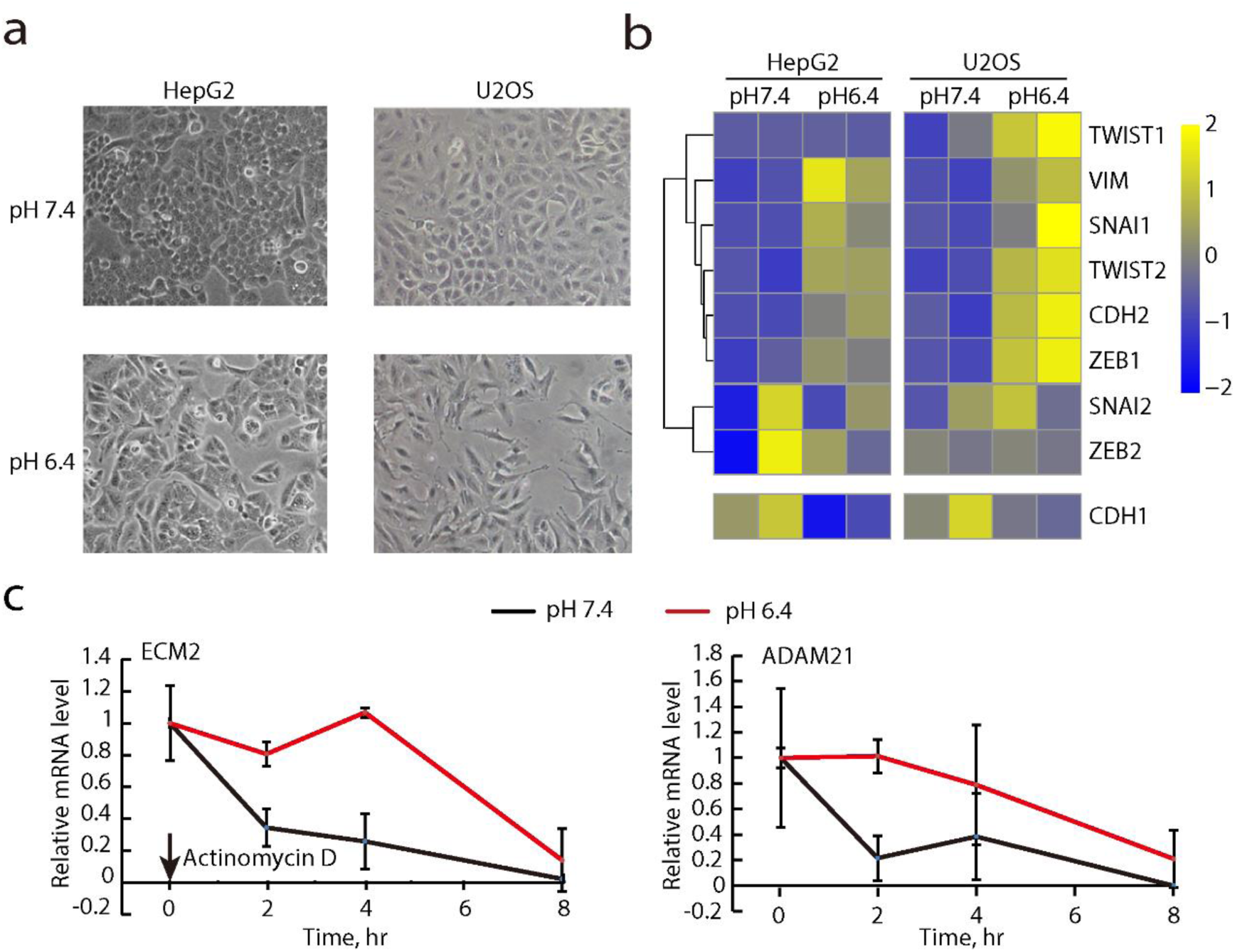
Acidic pH stabilizes ECM genes in cancer cells. (**a**) Morphology of human cancer cell lines HepG2 and U2OS at different pH; (**b**) Heatmap showing mRNA expression of epithelial (Cdh1) and mesenchymal markers in human cancer cell lines HepG2 and U2OS. (**c**) mRNA degradation plot of two representative ECM genes. The β-actin gene was used as a control. For each gene, data were normalized to U2OS cells cultured in each condition without actinomycin D treatment (0 hour). Shown are mean ± SD, n = 2. Actinomycin D was added 48 hr after low pH treatment, and cells were collected at 0, 2, 4, and 8 hr after actinomycin D addition for RNA-seq.

**Figure S7, Related to Figure 6.**
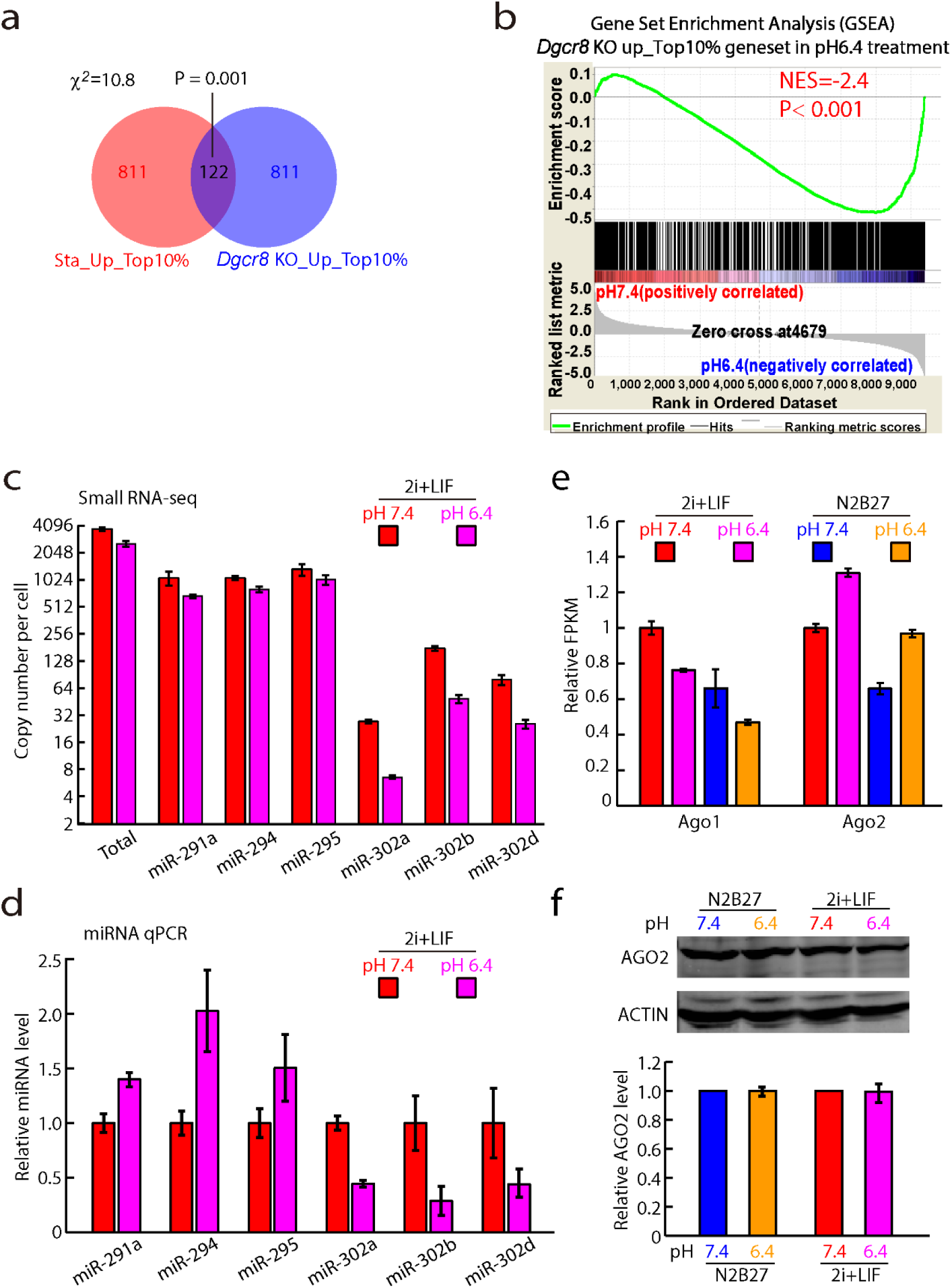
Low pH downregulates AGO1 protein expression and de-represses a subset of miRNA targets. (**a**) Venn diagrams showing the overlap between genes upregulated in Dgcr8 knockout mouse ESCs and genes with increased half-life in low pH treated mouse ESCs; (**b**) GSEA analysis showing the enrichment of upregulated genes in Dgcr8 knockout versus wild type ESCs in low pH treated ESCs; (**c**) Expression levels of miR-290/302 family of miRNAs in pH 6.4 and pH 7.4 ESCs from small RNA-seq. (**d**) Expression levels of miR-290/302 family of miRNAs in pH 6.4 and pH 7.4 ESCs from miRNA qPCR. Data were normalized to 7SK RNA and then to 2i+LIF pH 7.4 ESCs. Shown are mean± SD, n = 2. (**e**) Expression levels of Ago1 and Ago2 in different culture conditions from RNA-seq. Data were normalized to FPKM in 2i+LIF pH 7.4 ESCs. Shown are mean± SD, n = 2. (**f**) Western analysis AGO2 of mESCs in different culture conditions, The ACTIN was used as a loading control. Top panel: representative gel images. Bottom panel: quantification of AGO2 levels. Data were normalized to ACTIN then to pH 7.4 media. Shown are mean ± SD, n = 2.

